# Analysis of spatiotemporal specificity of small RNAs regulating hPSC differentiation and beyond

**DOI:** 10.1101/784819

**Authors:** Lu Li, Jin Feng Li, Dan Dan Cao, Vassilios Papadopoulos, Wai Yee Chan

**Author notes:** Corresponding author, (L.L.); (W.Y.C.).

## Abstract

We present a quantitative analysis of small RNA dynamics during the transition from hPSCs to the three germ layer lineages to identify spatiotemporal-specific small RNAs that may be involved in hPSC differentiation. To determine the degree of spatiotemporal specificity, we utilized two algorithms, namely normalized maximum timepoint specificity index (NMTSI) and across-tissue specificity index (ASI). NMTSI could identify spatiotemporal-specific small RNAs that go up or down at just one timepoint in a specific lineage. ASI could identify spatiotemporal-specific small RNAs that maintain high expression from intermediate timepoints to the terminal timepoint in a specific lineage. Beyond analyzing single small RNAs, we also quantified the spatiotemporal-specificity of microRNA families and observed their differential expression patterns in certain lineages. To clarify the regulatory effects of group miRNAs on cellular events during lineage differentiation, we performed a gene ontology (GO) analysis on the downstream targets of synergistically up-and downregulated microRNAs. To provide an integrated interface for researchers to access and browse our analysis results, we designed a web-based tool at https://keyminer.pythonanywhere.com/km/.

## Introduction

Human pluripotent stem cells (hPSCs) have emerged as a new model system for understanding the mechanism underlying human embryonic development ^1^. In addition, the functional cells derived from hPSCs have been considered as novel cell sources for replacement therapy and drug selection ^2–4^. Identification of critical members of different molecule classes regulating the hPSC differentiation process is essential for hPSC-based clinical applications that require a comprehensive understanding of both physiological and pathological mechanisms. hPSC differentiation is substantially regulated in both lineage and time, which is analogous to “spatiotemporal regulation” in human embryogenesis ^5, 6^. Therefore, the molecules that change in a spatiotemporal-specific manner may also control hPSC differentiation in both dimensions.

The profiling study of transcriptional and epigenetics dynamics during the differentiation of hPSCs, reported by Gifford *et al*, clarified the transcriptome changes, DNA methylation alterations, and chromatin modification dynamics during the formation of the three germ layers derived from hPSCs ^7^. In the same year, Xie *et al* profiled the transcriptome and epigenome of several cell states differentiated from hPSCs that represent key developmental decisions in the embryo ^8^. From these efforts, an integrated atlas of spatiotemporal dynamics of hPSC differentiation began to emerge. However, for the class of small RNAs, which is a critical player in guiding the differentiation of hPSCs ^9–14^, there is less information in this atlas. To date, the most comprehensive analysis of small RNA abundance in human tissues was performed by Ludwig *et al*, revealing the specific distribution of microRNAs (miRNAs) in mature tissues ^15^. For the three germ layers and their later cell derivatives corresponding to less mature states, the miRNA transcriptome (miRNAome) remains largely unexplored. In addition, most of the previous studies focused only on miRNAs while ignoring other small RNAs, including pre-miRNAs, snoRNAs, CDBox RNAs, H/ACA Box RNAs, and scaRNAs, which also could potentially regulate early embryogenesis ^16^.

Small RNAs are considered as master regulators of numerous transcription factors that directly control hPSC differentiation ^17, 18^. In general, when small RNAs regulate differentiation, they are either transiently upregulated in the intermediate state of differentiation or maintained with high expression to the mature state ^14, 19–21^. In our studies, if such temporal changes in expression appear in only one lineage, we defined them as spatiotemporal-specific small RNAs.

Recently, our group has profiled the expression dynamics of small RNAs during differentiation of human induced PSCs (hiPSCs) toward three key lineages (hepatic, nephric and neuronal) that are representative of the three germ layers ^22^. Using this dataset, we performed a hierarchical clustering analysis to reveal spatiotemporal-specific small RNAs. However, since this analysis tends to be biased against spatiotemporal-specific small RNAs with less change over time, this analysis provided only an incomplete list. Further, it was incapable of indicating the degree of spatiotemporal specificity of any of the hits. In this paper, we performed a quantitative analysis of the expression dynamics of small RNAs to determine their degrees of spatiotemporal specificity. In addition, our quantitative algorithms enabled the identification of small RNAs with unique expression patterns even their changes in expression might be small.

We developed two different methods to determine the spatiotemporal specificity. The first method used a normalized maximum timepoint specificity index (NMTSI) to identify changes in spatiotemporal-specific small RNA expression levels at just one timepoint in comparison to other timepoints. The second method used an across-tissue specificity index (ASI) to find spatiotemporal-specific small RNAs that are either specifically expressed at the terminal timepoint or maintained at high expression levels from an intermediate to the terminal timepoint.

Beyond the analysis of single small RNA, we further investigated the group behaviour of small RNAs, which is important for understanding how small RNAs contributes to differentiation when they cooperate with each other. We looked at the spatiotemporal specificity of miRNA families, which have been reported as clusters of small RNAs that share common targets and regulate signalling pathways synergistically ^23, 24^. To show the spatiotemporal specificity of miRNA families, we calculated mean ASI and number of spatiotemporal-specific miRNAs inside each family. Besides, we also analyzed the functions of spatiotemporal-specific miRNAs that are synergistically changed in expression. To clarify their regulatory effects on cellular events during lineage differentiation, we performed a gene ontology (GO) analysis on the downstream targets of co-upregulated or co-downregulated miRNAs.

To provide easy access to our dynamic small RNAs expression atlas, we implemented a web-based repository that includes all important analysis results. The information of small RNAs can be retrieved using their names as the key. This website is freely available at https://keyminer.pythonanywhere.com/km/.

## Results

### Identification of spatiotemporal-specific small RNAs with NMTSI

In our previous study, the profiling was performed on hepatocyte differentiation (HD), nephron progenitor differentiation (KD), and neural progenitor differentiation (ND) that were derived from the same hPSCs. Samples for profiling were collected at day 0, 3, 6, 10 of differentiation of the three lineages. In this study, we searched for the small RNAs that go up or down at just one timepoint in a specific lineage.

We first evaluated the degree of temporal specificity of small RNAs. Briefly, we used timepoint specificity index (TSI) as a graded scalar to measure the specificity of expression of a small RNA with respect to different timepoints ^15, 25, 26^. TSI of single small RNAs in HD, KD, and ND were calculated separately and summarized in Table S1 (column C-E). Since the normalization may affect the results ^15^, we performed all analyses for TSI on raw data using raw expression intensity values (expression values). TSI has a range of 0 to 1, representing the expression dynamics of any small RNA ranging from ubiquitous expression at all timepoints (0) to specific expression at only one timepoint (1). In HD, 78.6% of all small RNAs showed an intermediate TSI value (Fig. 1A). Similarly, 79.3% of all small RNAs and 79.2% of all small RNAs showed an intermediate TSI value (0.1 to 0.6) in KD and ND, respectively (Fig. 1B, C), suggesting that most small RNAs change moderately during lineage differentiation.

**Figure 1.**
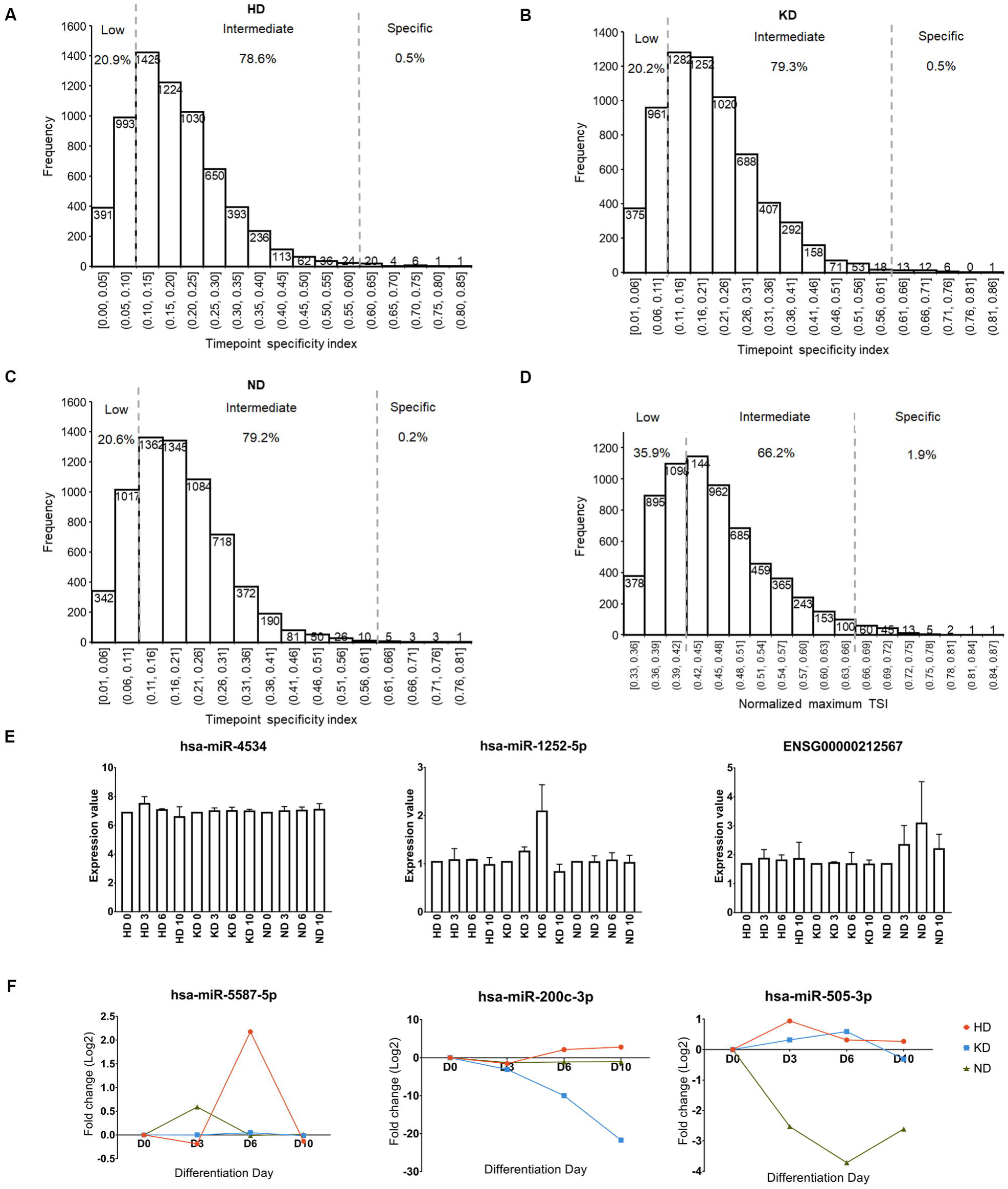
Characterization of spatiotemporal-specific small RNA identified by NMTSI and DE. (A-C) Histogram plot for the frequency of timepoint specificity index (TSI) of small RNAs detected in hepatocyte differentiation (HD), nephron progenitor differentiation (KD), and neural progenitor differentiation (ND). The vertical dotted lines correspond to the threshold proposed for defining low expressed (< 0.11) and specifically expressed small RNAs (> 0.61). (D) Normalized maximum TSI (NMTSI) distribution of small RNAs detected in hPSC differentiation. The vertical dotted lines correspond to the threshold proposed for defining low expressed (< 0.42) and specifically expressed small RNAs (> 0.66) of spatiotemporal-specific candidates. (E) Bar plot of expression values (mean ± SD) of spatiotemporal-specific candidate small RNAs with the highest NMTSI value in HD, KD, and ND, respectively. (F) Dynamic expression patterns of spatiotemporal-specific small RNAs with the highest NMTSI value in HD, KD, and ND, respectively.

Next, we established the spatiotemporal specificity of small RNAs as the lineage with the highest TSI among HD, KD, and ND (Table S1, column F). We also estimated the degree of spatiotemporal specificity by the NMTSI value, which was calculated by comparing the “weight” of TSI obtained from HD, KD, and ND (Table S1, column G). NMTSI values ranged from 0.33 to 0.86 (Fig. 1D), with values close to 1 stand for small RNAs uniquely upregulated in only one lineage and values close to 0.33 represent small RNAs either upregulated or unchanged together in all lineages. In total, 35.9% of small RNAs showed a low NMTSI < 0.42 and 62.2% of small RNAs showed an intermediate NMTSI (0.43 to 0.66) (Fig. 1D). 126 small RNAs showed a high NMTSI (0.67 to 0.88) (Fig. 1B), suggesting a strong spatiotemporal specificity in hPSC differentiation.

By ranking NMTSI values from the highest to the lowest in Table S1, the top 100 small RNAs were revealed (NMTSI > 0.67, labelled in the blue background in Table S1). Particularly, *hsa-miR-4534*, *hsa-miR-1252-5p*, and *ENSG00000212567* (snoRNA) were the most spatiotemporal-specific small RNAs in HD, KD, and ND, respectively. Figure 1E showed that KD-specific *miR-1252-5p* and ND-specific *ENSG00000212567* were significantly changed at day 6 of KD (KD 6) and ND 6, suggesting that NMTSI is quite accurate in identifying spatiotemporal-specific candidate small RNAs. Different from obvious changes, the upregulation of HD-specific *miR-4534* at HD 3 was mild (Fig. 1E), indicating that NMTSI is sensitive in identifying spatiotemporal-specific candidate small RNAs despite their small changes in expression.

A concern remains that a small change in expression may be due to noise presented in low intensities, for which a further filtration with statistical significance should correct. For each small RNA, we processed the raw expression value to obtain the fold-change value and false discovery rate (FDR)-value when comparing the raw expression value between any of two timepoints. Thereafter, we set a cut-off of FDR < 0.05 with any fold-change (differential expression, DE) to find those small RNAs with a significant change in expression. By filtering candidate spatiotemporal-specific small RNAs obtained by NMTSI with a cut-off of DE, we got the final list of spatiotemporal-specific small RNAs (Table S2). In total, 330 HD-, 123 KD-, and 677 ND-specific small RNAs were identified. Their NMTSI values, spatiotemporal specificity, fold-change values, and FDR*-*value were summarized in Table S2.

By ranking NMTSI values from maximum to minimum in the final list (Table S2, column D), we observed that *hsa-miR-5587-5p*, *hsa-miR-200c-3p*, and *hsa-miR-505-3p* showed the highest NMTSI value in HD, KD, and ND, respectively. Their normalized fold-change values were plotted in Figure 1F. All of them showed a timepoint-specific expression in just one lineage, supporting the accuracy of the combination of NMTSI and DE in the identification of spatiotemporal-specific small RNAs.

### Identification of spatiotemporal-specific small RNAs with ASI

In the analysis above, we used NMTSI to identify those small RNAs specifically going up or down at a single timepoint. However, some cell-fate determinants have been found to be highly expressed at more than one timepoint ^14, 27, 28^. To identify these small RNAs, we used ASI to measure their degree of specificity at the terminal timepoint of different lineages.

In the dataset we studied (Data citation 1), day 10 of HD-, KD-and ND-cells correspond to three immature tissues, namely, immature hepatocytes (IH), metanephric mesenchyme (MM), and neural progenitor cells (NPC). For each small RNA, we calculated its ASI using the expression values of IH, MM and NPC. ASI values of all small RNAs were summarized in Table S3 (column F). ASI has a range of 0-1, indicating the distribution of small RNAs from ubiquitous expression (0) to specific expression (1) among various tissues. In total, 83.9% of small RNAs showed an intermediate ASI (0.1-0.6) (Fig. 2A). 110 small RNAs (1.7%) showed a high ASI (Fig. 2A), suggesting a strong spatial specificity in hPSC differentiation. In parallel, the spatial specificity of each small RNA (column G) was established as the lineage with the highest expression value at day 10.

**Figure 2.**
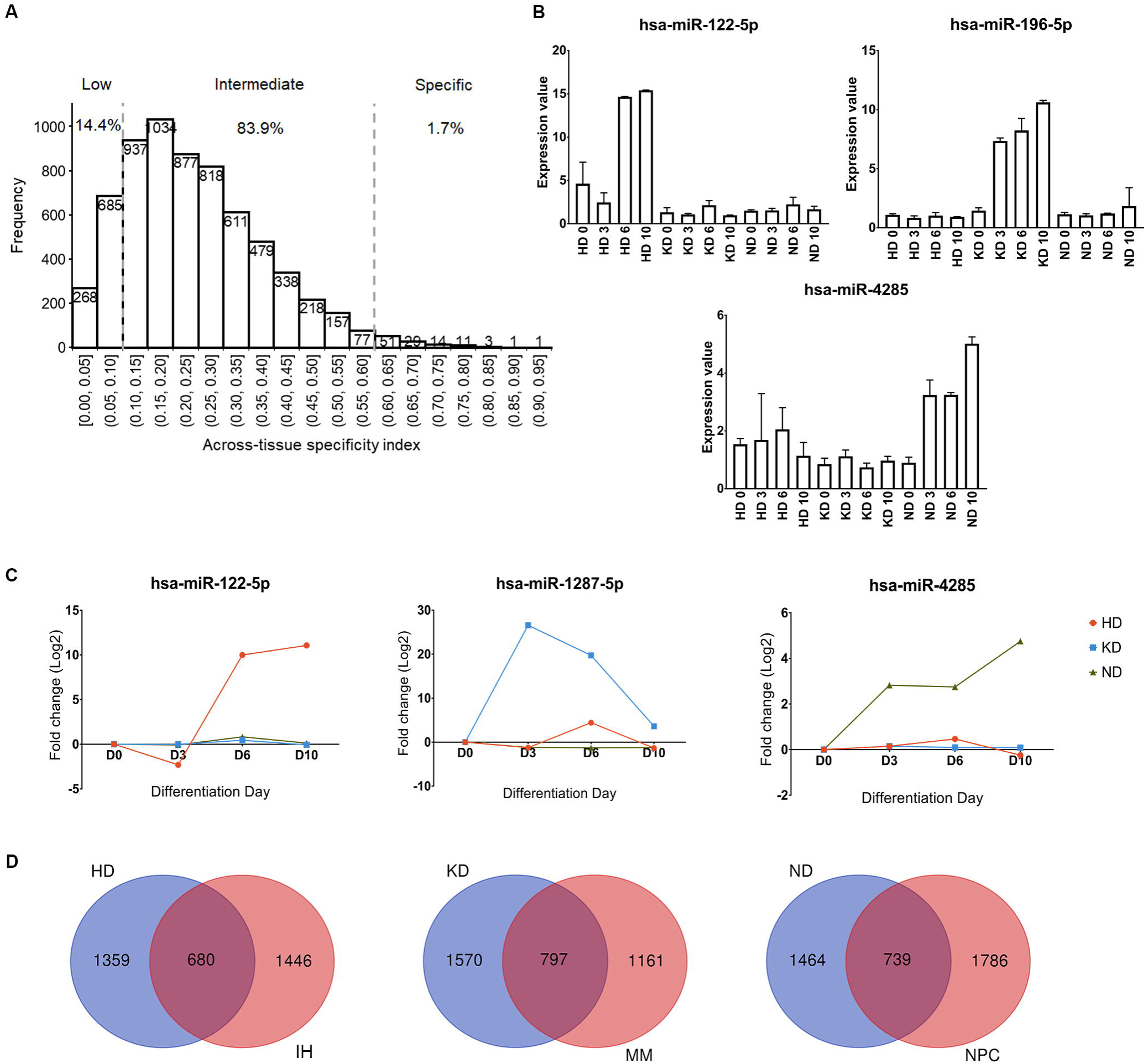
Characterization of spatiotemporal-specific small RNA identified by ASI and DE. (A) Across-tissue specificity index (ASI) distribution of small RNAs detected in hPSC differentiation. The vertical dotted lines correspond to the threshold proposed for defining low expressed (< 0.10) and specifically expressed small RNAs (> 0.60) in tissue-specific candidates. (B) Bar plot of expression values (mean ± SD) of tissue-specific candidate small RNAs with the highest ASI value in immature hepatocytes (IH), metanephric mesenchyme (MM), and neural progenitors (NPC), respectively. (C) Dynamic expression patterns of spatiotemporal-specific small RNAs with the highest ASI value in IH, MM, and NPC, respectively. (D) Venn diagram of spatiotemporal-specific candidate small RNAs indicated by NMTSI analysis and spatiotemporal-specific candidate small RNAs indicated by ASI analysis in HD, KD, and ND, respectively.

Sorting ASI values from maximum to minimum in Table S3, we found that *hsa-miR-122-5p* showed the highest ASI value (0.91). Since it is expressed the highest in IH among the three tissues, it is an IH-specific small RNA. Remarkably, its high level of expression is maintained from HD 6 to 10 (Fig. 2B), indicating that the capacity of ASI to identify tissue-specific small RNAs is not limited by the expression duration of small RNAs. Similarly, by ranking ASI values in Table S3, we found *hsa-miR-196a-5p* (ASI=0.87) and *hsa-miR-4285* (ASI=0.79) with the highest ASI in MM and NPC, respectively, are highly expressed at more than two timepoints (Fig. 2B).

After establishing the spatial specificity of small RNAs, we determined the small RNAs, which also showed a temporal specificity. By filtering spatio-specific small RNAs obtained by ASI with a cut-off of DE (FDR-value < 0.05), the spatio-specific small RNAs with a temporal DE detected between any of two timepoints were revealed. In total, 195 IH-, 104 MM-, and 1019 NPC-specific small RNAs were identified as spatiotemporal-specific small RNAs. Their ASI values, spatiotemporal specificity, normalized fold-change values, and FDR*-*value were summarized in Table S4.

By ranking the ASI from the highest to the lowest in Table S4 (column D), *hsa-miR-122-5p*, *hsa-miR-1287-5p*, and *hsa-miR-4285* showed the highest ASI in IH, MM, and NPC, respectively. Their normalized fold changes were plotted in Figure 2C. Not surprisingly, they are all sustained high expression from intermediate timepoints to the terminal timepoint.

Notably, the lists of spatiotemporal-specific small RNAs identified by NMTSI and ASI partially overlap (Fig. 2D). It is probably due to ASI, which only counts the enrichment of small RNAs at day 10, leading to an inclusion of both small RNAs specifically expressed at day 10 and small RNAs with sustained high expression from intermediate timepoints.

### Spatiotemporal specificity of miRNA families

Beyond looking into the spatiotemporal specificity of single small RNAs, we were also interested in the spatiotemporal specificity of small RNA clusters, e.g. miRNA families. We explored the lineage in which single miRNA families showing specific expression dynamics from the pluripotent state (day 0) to the differentiating states (day 10). To determine the degree of the spatiotemporal specificity of miRNA families, we calculated the mean ASI for spatiotemporal-specific miRNAs inside each miRNA family. To display the distribution of miRNA families, the number of detected family members in IH, MM, and NPC was counted separately.

In total, 115 IH-, 69 MM-, and 553 NPC-specific miRNAs were extracted from Table S4 based on the type of small RNAs (column C). According to previous papers ^15, 25^, we focused on the miRNA families containing at least five known mature miRNAs. Therefore, 37 out of 589 miRNA families extracted from the miRbase V21 were analyzed ^29^.

For families with more than 3 members detected, mir-302 family, mir-25 family, and mir-506 family showed the highest mean ASI in IH, MM, and NPC, respectively (Fig. 3A-C, indicated by orange circles), suggesting a strong spatiotemporal specificity in lineage differentiation. Moreover, since mir-500 family, mir-10 family and, mir-515 family have the greatest number of detected family members in IH, MM, and, NPC, respectively, they showed the most biased distribution in lineage differentiation among all families (Fig. 3 A-C, indicated by blue circles). The specificity and distribution of miRNA family members in three lineages were summarized in Table S5.

**Figure 3.**
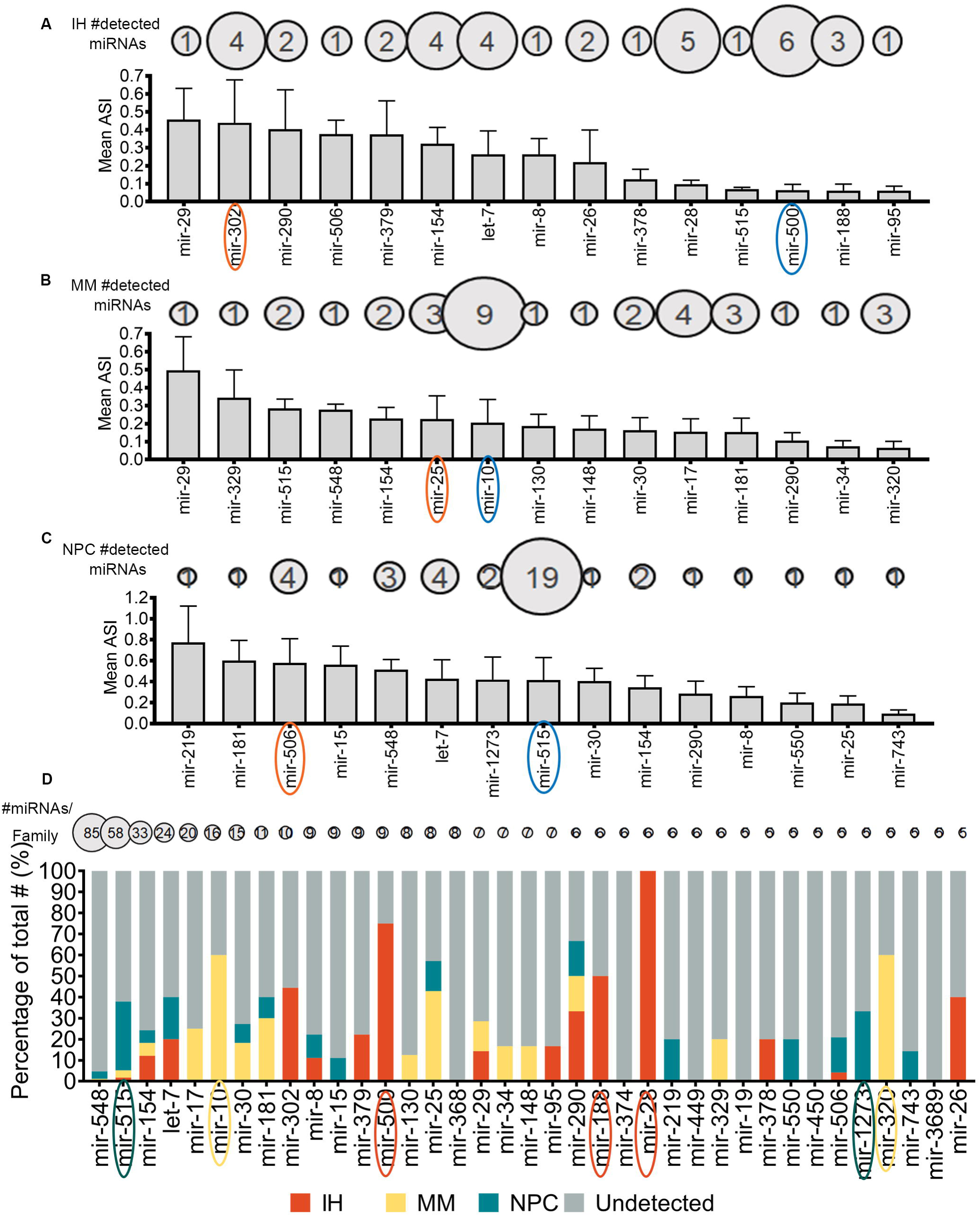
Spatiotemporal specificity of miRNA families. Average of ASI in different miRNA families in (A) IH, (B) MM, and (C) NPC. For each miRNA family with at least five known members, the mean and standard deviation of detected family members ASI in a certain lineage is presented as a bar plot. Families are sorted with decreasing average ASI from left to right. The number of detected family members is shown above columns with balloons, representing the detected size. MiRNA families with the highest mean ASI in IH, MM, and NPC are indicated by orange circles. MiRNA families with the biggest number of spatiotemporal-specific family members in IH, MM and NPC are indicated by blue circles. (D) Inside each family, the number of IH-, MM-, NPC-, and undetected family members were compared, and the proportions presented as a percentage stacked bar plot. The number of total family members is shown above columns with balloons, representing the family size. MiRNA families showing a particularly high specificity in IH, MM, and NPC are indicated by red, yellow, and green circles, respectively.

We then investigated the distribution of 37 families in lineage differentiation by estimating the percentage of spatiotemporal-specific family members inside a family in each lineage. Mir-28, mir-500, and mir-188 family showed a particularly high specificity in IH (≥ 50 % of family members were IH-specific) (Fig. 3D, indicated by red circles). mir-320 and mir-10 family showed a particularly high specificity in MM (≥ 50 % of family members were MM-specific) (Fig. 3D, indicated by yellow circles). However, in ND, only mir-1273 and mir-515 family showed a slight NPC-specificity (≥ 30% of family members were NPC-specific) (Fig. 3D, indicated by a blue circle), implicating that the formation of NPC is fine-tuned by a complicated miRNA regulatory system rather individual miRNA families.

If considering both a high average of ASI values (Fig. 3A-C) and a great percentage of spatiotemporal-specific members (Fig. 3D), mir-500 family, mir-10 family, and mir-515 family are the most spatiotemporal-specific miRNA families.

### Functional analysis of spatiotemporal-specific miRNAs

We further interpret the cellular events elicited by small RNAs expression dynamics. Given that the target genes of mature miRNAs are well-studied, we focused on the spatiotemporal miRNAs. Based on the list of spatiotemporal-specific miRNAs obtained by ASI (Table S4), we classified these miRNAs into six groups, corresponding to the miRNAs being upregulated at HD 10, KD 10 and ND 10 (fold change >1 and FDR < 0.05 when compared day 10 to day 0) or miRNAs being downregulated at HD 10, KD 10 and ND 10 (fold change < −1 and FDR < 0.05 when compared day 10 to day 0). The six groups are listed in Table S6 (column A, C, E, G, I, K). We then searched for target genes for each group of miRNAs using miRTarBase database. Their targets were summarized in Table S6 (column B, D, F, H, J, L).

Next, we performed a GO analysis on each group of targets using the web tool PANTHER version 14.1 (http://www.pantherdb.org/tools/)^30^. Statistical overrepresentation test terms under the “Gene List Analysis” function with FDR < 0.05 were considered significantly enriched. Fold enrichment was used as the ranking criteria (Table S7, column F). Consequently, we observed that different GO terms were associated with up-and downregulated miRNAs in these lineages (Table S7, column A). For example, in ND, the targets of upregulated miRNAs were enriched in the pathways related to brain development and neural tube formation (Fig. 4A); however, the targets of downregulated miRNAs were associated with eye development and noncoding RNA (ncRNA)-mediated translation process (Fig. 4B). The top 10 enriched GO terms for each group were clustered manually into biologically related topics (Fig. 4A, B). All GO terms associated with six group of miRNAs were summarized in Table S7.

**Figure 4.**
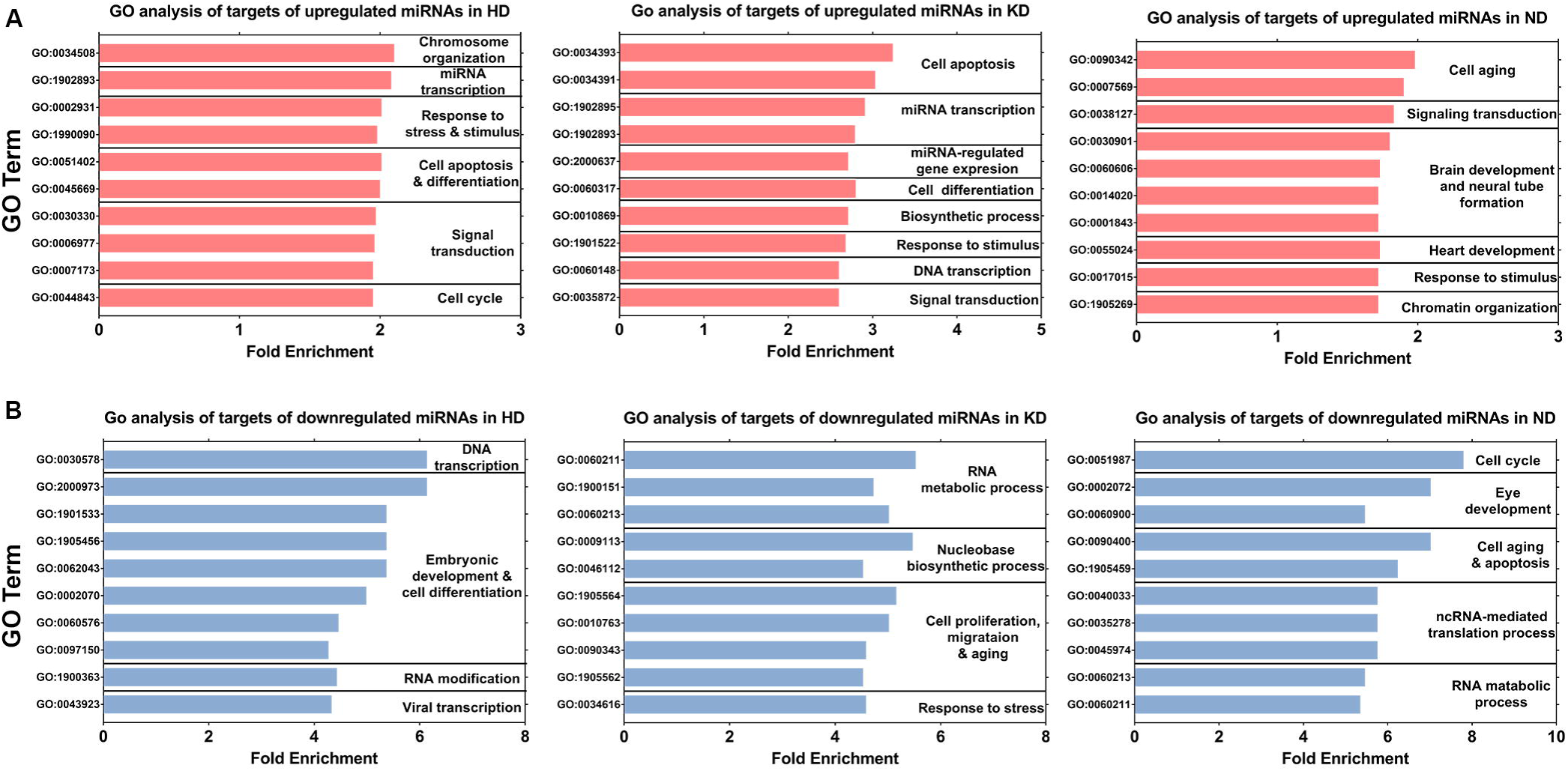
Identification of functions of spatiotemporal-specific small RNAs during hPSC differentiation. Gene ontology (GO) analysis for downstream targets of (A) upregulated miRNAs and (B) downregulated miRNAs. The 10 most highly enriched GO biological process terms are manually clustered into related topics for HD, KD and ND.

### The search engine for small RNAs of interests

In order to share our experimental results with the scientific community at large, we developed a search engine using Django that can quickly retrieve information with regard to any specific small RNA. We have deployed this search engine on a website, such that visitors can type in the name of a small RNA to retrieve all related information. The search engine is available at: https://keyminer.pythonanywhere.com/km/.

When a visitor types in the name of a small RNA of interest (e.g., *hsa-miR-181b-5p*), Django will pass this name as a text to our server. On receiving this text, our server will look for the row that matches this text and return all data in this row back to the browser for visualization. The front-end of the browser displays the returned data in a pre-defined HTML table. The results will be shown as four separate parts on the website (Fig. 5). The first part is the basic information of the searching small RNA, including probe ID (e.g., MIMAT0000257_st), name, NMTSI, ASI, potential spatiotemporal specificity indicated by NMTSI (HD, KD, or ND), potential spatiotemporal specificity indicated by ASI (IH, MM, or NPC). The second part is the normalized fold change of the specified small RNA during differentiation of the three lineages. The third part is the actual spatiotemporal specificity of a specified small RNA. If the specified small RNA has no actual spatiotemporal specificity after a filtration with DE, the result will be shown as a “#N/A”. The fourth part is the miRNA family distribution, if a spatiotemporal-specific miRNA is specified and only if this miRNA belongs to a miRNA family containing at least five members, the miRNA family name (e.g., mir-181) and the family distribution figure will be shown. Our website is able to replace cells in these HTML tables without reloading whole webpages. An example of searching for a small RNA (*hsa-miR-181b-5p*) is shown in Figure 5.

**Figure 5.**
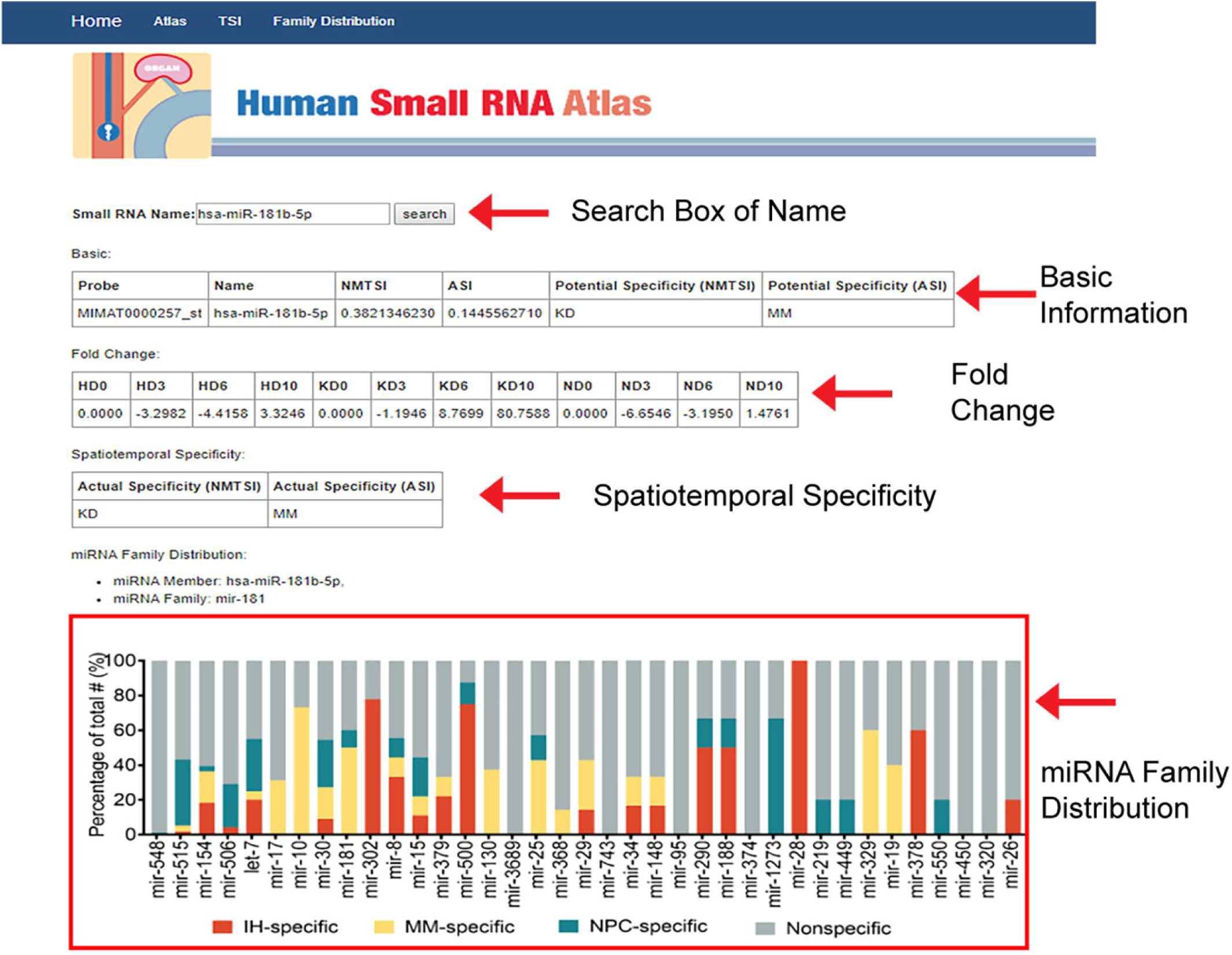
The search engine for small RNAs of interests. The small RNA database entry for *hsa-miR-181b-5p*. Four sections of the page display from top to bottom: the basic information, detected fold-changes in HD, KD and ND (day 0, 3, 6, 10), spatiotemporal specificity (indicated by NMTSI and ASI, respectively), and miRNA family distribution. The actual spatiotemporal specificity of *hsa-miR-181b-5p* (MM, identified by ASI and DE) indicates that it is located at the yellow stack of the family mir-181 bar.

## Discussion

Since small RNAs have emerged as crucial cell fate determinants, there is an increasing need for the identification of small RNAs directing hPSC differentiation ^31^. Tissue-specific small RNAs have been implicated in the regulation of lineage formation and in the maintenance of tissue properties ^14, 19–21, 32^, whereas the small RNAs that determine early lineage differentiation remains largely unknown. Although there are ample studies investigating differentially expressed small RNAs during lineage differentiation, most of them profile only the expression dynamics in one lineage. Without comparisons among multiple lineages, it is difficult to determine whether the expression patterns are general in the reduction of stemness or unique in the differentiation of certain lineages. Additionally, due to the variable genetic background of hPSC donors and different profiling platforms, the results of these studies are difficult to integrate, limiting their applications in the comparison of small RNA dynamics. While our previous study profiled small RNA dynamics of multilineage differentiation derived from the same hPSCs avoided these concerns (Data citation 1), it still had limitations on fully identifying all spatiotemporal small RNAs involved in early differentiation.

Using our new analytical approach, we found more spatiotemporal-specific small RNAs (282 vs 1615). The increment in identified numbers are due to three main reasons: First, we adopted new analysis algorithms, namely NMTSI and ASI, as compared to hierarchical clustering algorithms used previously. NMTSI and ASI are quantitative scalar measures for the specificity of expression of small RNAs ^15^. Their capacity to quantify spatiotemporal specificity allowed a more thorough investigation of spatiotemporal-specific small RNAs with either great or small expression changes when sorting the indices from the highest to the lowest. Secondly, using ASI we could identify small RNAs sustained high expression at more than one timepoint that are complementary to NMTSI identified small RNAs that are highly expressed at only one timepoint. This classification ensured more comprehensive coverage of spatiotemporal-specific small RNAs. Thirdly, we used absolute fold change > 1 instead of 2 as the criteria when filtering spatiotemporal-specific candidate small RNAs, by which the number of spatiotemporal-specific small RNAs was increased.

After the DE filtration, we observed an alteration in the list of the most spatiotemporal-specific small RNAs (Fig. 1E), in which *hsa-miR-4534*, *-1252-5p*, and *ENSG00000212567* were filtered out. Instead, *hsa-miR-5587-5p*, *-200c-3p*, and *-505-3p* became the most timepoint-specific small RNAs (Fig. 1F). *hsa-miR-4534* was filtered out due to a small change in expression during HD (Fig. 1E). *hsa-miR-1252-5p* and *ENSG00000212567* were excluded due to a non-significant difference (FDR-value > 0.05), as implicated by the large bias of expression values between biological duplicates (Fig. 1E). Therefore, the DE filtration is necessary to exclude false positive results.

In the analysis of ASI and DE*, hsa-miR-122-5p*, *-1287-5p* and *-4285* have been identified as the most spatiotemporal-specific small RNAs (Fig. 2B, C). Consistent with existing studies, *hsa-miR-122-5p* is specifically correlated with hepatocyte formation ^33–35^, while *hsa-miR-1287-5p* and *-4285* might be potential regulators of MM and NPC formation that require further studies.

Beyond the identification of single spatiotemporal-specific small RNAs, we also investigated the spatiotemporal specificity of miRNA families. For miRNA family members specifically distributed in single lineages, we calculated both mean ASI and the number of family members inside each family with respect to individual lineages to show their spatiotemporal specificity^15^. We found that the neural progenitor-enriched let-7 family ranking 6th in mean ASI and 2nd in the number of family members (Fig. 3C), which suggested a strong spatiotemporal specificity in ND. This result is in line with previous studies that report an enrichment of let-7 family in neural progenitors^36, 37^.

Moreover, we found that several families presenting a spatiotemporal-specific distribution were previously unknown, such as IH-specific mir-302 family, MM-specific mir-10 family, and NPC-specific mir-515 family (Fig. 3A-C). Novel identification of these spatiotemporal-specific families shall aid in understanding how a miRNA family influences on lineage specification. Furthermore, the co-expression pattern of family members narrows down the range of downstream targets that helps to efficiently unmask critical cellular events accompanying hPSC differentiation ^38^. Of note, given their co-expression patterns and redundant functions in regulating signalling pathways, it may be better to target the whole family instead of single miRNAs when studying the effects of miRNAs on lineage differentiation ^39^.

To further clarify the cellular events associated with spatiotemporal-specific small RNAs, we focused on groups of miRNAs that are synergistically upregulated or downregulated. The GO analysis of such targets revealed that biological processes were differentially associated with individual lineages. For examples, embryonic development, RNA metabolic process, and brain development are apparently correlated with HD, KD, and ND, respectively (Fig. 4A, B). Some of the cellular events are more likely responses to “stimuli” (growth factors and chemicals) added to the induction medium, whereas others may be triggered by the master factors (miRNAs and transcription factors) changed during differentiation. Notably, we used fold enrichment instead of the *P* value as the ranking criterion for biological processes, since we observed that the ones with a large number of expected genes (e.g. cellular metabolic process, cellular process) were always ranked in the top when considering the *P*-value. However, sorting fold enrichment, which reflects the ratio of overrepresented number of genes in the uploading list compared to the expected number of genes in the reference list ^30^, allows the identification of other important biological processes to rank top despite small expected numbers of genes.

Taken together, our analysis filled the void with respect to small RNA expression dynamics in the human atlas for hPSC differentiation. Our results can be used as informative clues for investigating spatiotemporal-specific small RNAs and their roles in key decisions in human developmental processes. Meanwhile, the analysis framework can serve as a template for the comparisons of dynamic spatiotemporal transcriptome changes during diverse multilineage differentiation.

## Materials and Methods

### Analysis of Small RNA expression in hPSCs and derived lineages

Small RNAs microarray data of hPSC differentiating into three lineages have been published previously (Data citation 1) ^22, 40–42^. In brief, hPSCs were induced into representative lineages according to previously established protocols (hepatic, nephric and neuronal) ^40–42^. RNA was extracted at four different timepoints (day 0, 3, 6, 10) for each lineage using the RecoverAll™ Total Nucleic Acid Isolation Kit for FFPE (Thermo Fisher Scientific) and subjected to the microarray-based small RNA expression analysis using the Affymetrix miRNA 4.0 platform (Thermo Fisher Scientific) as described ^22^. This dataset has been deposited in the gene expression omnibus (GEO) repository under the accession number GSE97952 (Data citation 1). The raw expression intensity values (expression values) were extracted using Partek® Genomics Suite® platform. To show human-specific information, human-specific probes were specifically selected.

### TSI, NMTSI, and ASI

To evaluate the variability of temporal expression patterns inside each lineage, we calculated a TSI for single small RNAs as described before ^25^. This specificity index is a quantitative measurement for the expression specificity of small RNAs with re tard to different timepoints. TSI has a range of 0 to 1, with values approximate to 0 representing small RNAs that remained unchanged during differentiation and values approximate to 1 representing small RNAs that were expressed at only one timepoint. Considering that the fluctuation of expression values of the start timepoint (HD 0, KD 0, ND 0) have potential effects on the comparison of TSI between lineages, we used the average value of day 0 from three lineages (mean of HD 0, KD 0, and ND 0) as the expression value for HD 0, KD 0, and ND 0. The TSI for a small RNA *j* is calculated as

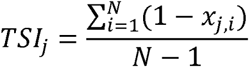

where N corresponds to the total number of timepoints measured and *x_j,i_*is the expression value of timepoint *i* normalized by the maximum expression value of any timepoint for small RNA *j*.

To further identify the small RNAs that are changed in a timepoint-specific manner in only one lineage (spatiotemporal-specific small RNAs), we developed the Normalized Maximum TSI (NMTSI) based on TSI values. NMTSI values range from 0.33 to 1. The small RNA with NMTSI value 0.33 is either upregulated or unchanged over all lineages, while the small RNA with NMTSI value 1 is upregulated in only one lineage. The NMTSI for a small RNA *j* is calculated as

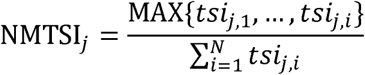

where N corresponds to the total number of lineage measured and *tsi_j,i_* is the TSI value of lineage *i* for small RNA *j*. Specifically, HD, KD, and ND were assigned lineage 1, lineage 2, and lineage 3, respectively, in this paper.

To identify spatiotemporal-specific small RNAs highly expressed at the terminal timepoint (HD 10, KD 10, and ND 10), we calculated an ASI for each small RNAs analogous to the tissue specificity index ‘τ’ originally developed for mRNA ^26^. Compared with TSI, ASI considers the variability of small RNA expression patterns between terminal timepoints of different lineages. The ASI for a small RNA *j* is calculated as

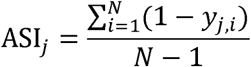

where N corresponds to the total number of lineages measured and *y_j, i_* is the expression value at day 10 of lineage *i* normalized by the maximum expression value at day 10 of any lineage for small RNA *j*.

### Identification of real spatiotemporal-specific small RNAs

To identify real spatiotemporal-specific small RNAs from the candidate list that was generated via NMTSI analysis, we performed a filtration with a differential expression (DE). To analyze DE patterns of small RNAs during lineage specification, we compared the small RNA transcriptome between any two timepoints inside a lineage. A workflow for miRNA microarray analysis launched at Partek® Genomics Suite® was used. Specifically, HD-, KD-, and ND-specific candidate small RNAs were filtered with a cut-off of DE (absolute fold-change value of > 1 and post hoc test FDR-value < 0.05).

Similarly, to identify real spatiotemporal-specific small RNA from the candidate list that was generated via ASI analysis, we applied a filtration with DE. Particularly, we filtered IH-, MM-, NPC-specific candidate small RNAs with a cut-off of DE (absolute fold-change value of > 1 and post hoc test FDR-value < 0.05).

### Expression of miRNA families

To evaluate the spatiotemporal specificity of miRNA families, we selected miRNA families containing at least five mature miRNAs from the miRbase V21 (ftp://mirbase.org/pub/mirbase/21/; catalogued as miFAM.dat.gz or miFAM.dat.zip; accessed on 06/26/2018). For each miRNA precursor all mature forms are considered as family members. Replicated mature miRNAs coming from different precursors were only counted once. Average of ASI and quantities of spatiotemporal-specific miRNA family members (identified by ASI and DE analysis) inside families with respect to the terminal timepoint of different lineages (IH, MM, and NPC) were calculated.

### *In silico* identification of target genes of miRNAs

Known human miRNA-target interactions were downloaded from the miRTarBase database (http://mirtarbase.mbc.nctu.edu.tw/php/download.php; catalogued as hsa_MTI.xlsx; accessed on 04/02/2019). Spatiotemporal-specific miRNAs (identified by ASI and DE analysis) in each lineage were classified into upregulate groups (fold change > 1 when compared day 10 to day 0) and downregulated groups (fold change < −1 when compared day 10 to day 0). Downstream target genes of each group of miRNAs were retrieved taking miRNA names as the key.

### GO enrichment analysis

The GO enrichment analysis for target genes was performed using the PANTHER version 14.1 (http://www.pantherdb.org/tools/). Statistical overrepresentation test with subterm “GO biological process complete” under the “Gene List Analysis” function with FDR < 0.05 were considered significantly enriched. Top 10 enriched GO terms (by fold enrichment) from each category were clustered manually into biologically related topics.

### The development of a search engine

We adopted Python and used the Django framework in developing our website. To organize the data in a structured way, we defined a Django model and associated this model with a database table. We adopted Sqlite as our database management system, which hosts this table and serializes its data into a single file. All of our data was inserted into Sqlite automatically in batches by a shell script. If there is more data in the future, we can also insert them with this shell script or by accessing the Django admin webpage.

Our Sqlite table contains 21 columns corresponding to the 21 aspects of a probed small RNA. These 21 aspects include the probe ID, small RNA name, NMTSI, ASI, potential spatiotemporal specificity (NMTSI), potential spatiotemporal specificity (ASI), fold-change of HD 0, HD 3, HD 6, HD 10, KD 0, KD 3, KD 6, KD 10, ND 0, ND 3, ND 6, ND 10, spatiotemporal specificity (NMTSI), spatiotemporal specificity (ASI), and miRNA family distributions. A row in this table uniquely records results for one probe. We assigned a probe ID rather than the small name as the primary key to index a row, since a small RNA may have multiple probes and thus take up multiple rows.

## Data Availability Statement

The data that support the findings of this study are openly available in Figshare at 10.6084/m9.figshare.9911918. The following have been uploaded to Figshare: expression values of all small RNAs; TSI values of small RNAs in HD, KD, and ND; NMTSI values of small RNAs for the calculation of NMTSI distribution; expression values of the most potential HD-, KD-, and ND-specific small RNAs; fold change of the most HD-, KD-, and ND-specific small RNAs; ASI values of small RNAs for the calculation of ASI distribution; expression values of the most potential IH-, MM-, and NPC-specific small RNAs; fold change of the most IH-, MM-, and NPC-specific small RNAs; average and SD of ASI of miRNA families in three lineages; distribution of 37 miRNA families in three lineages; and summary of top 10 GO terms related with spatiotemporal-specific small RNAs.

## Code Availability Statement

The scripts used to design the search engine are available from GitHub: https://github.com/keyminer/hsra.

## Data Citations

1. Li, L., Chan, W. Y. & Cheung, H. H. *Gene expression Omnibus* GSE97952 (2018).

## Author Contributions

L.L., and W.Y.C. conceived the project. L.L., J.F.L., and D.D.C. designed the experiments. L.L., and J.F.L. conducted the experiments. L.L., and J.F.L. analyzed the data and wrote the manuscript. W.Y.C. and V.P. co-wrote and edited the manuscript.

## Acknowledgements

This work was supported in part by the CUHK VC One-Off Discretionary Fund (Project 4930732) and the One-Off Funding for Joint Lab/Research Collaboration (Project 3132966) provided to the CUHK-CAS GIBH Joint Laboratory on Stem Cell and Regenerative Medicine, the Lo Kwee-Seong Biomedical Research Fund, and Project Stem Cell Therapy of Liver Diseases: An Investigation (2015CB964700), 973 Scheme, Ministry of Science and Technology, China.

## Table Legends

**Table S1. TSI, NMTSI and potential spatiotemporal specificity of small RNAs.** Summary of timepoint specificity index (TSI), normalized maximum TSI (NMTSI), and spatiotemporal specificity of 6609 small RNAs that are observed with microarrays. Column A and B show the ID of probes and corresponding name of small RNAs. The degree of temporal specificity of small RNAs is evaluated by TSI, the value of which in hepatocyte differentiation (HD), nephron progenitor differentiation (KD), and neural progenitor differentiation (ND) is shown in column C, D, and E, respectively. The spatiotemporal specificity of a small RNA is established as the lineage with the highest TSI among HD, KD, and ND (column F). The degree of spatiotemporal specificity of small RNAs is estimated by NMTSI values (column G) that is calculated using TSI values of HD, KD, and ND. Small RNAs are sorted with decreasing NMTSI values from top to bottom. Top 100 small RNAs with high NMTSI values (> 0.673) are indicated by the blue background.

**Table S2. Spatiotemporal-specific small RNA indicated by NMTSI and DE.** Listing of 1130 real spatiotemporal-specific small RNAs with both spatiotemporal specificity (from NMTSI analysis) and DE (absolute fold-change > 1 and post hoc test FDR-value < 0.05 between any of two timepoints). Column A to E show the probe ID, small RNAs name, small RNA type, spatiotemporal specificity, and NMTSI. Results of one-way ANOVA *P*-value, fold-change value and post hoc test FDR*-*value are generated from the comparison of expression values between any of two timepoints. These results are listed from column F to DN. HD-specific small RNAs (yellow background), KD-specific small RNAs (blue background), and ND-specific small RNAs (red background) are grouped together according to their spatiotemporal specificity.

**Table S3. ASI and spatial specificity of small RNAs.** Summary of mean expression values of the terminal timepoint of the three lineages (HD, KD, and ND), across-tissue specificity index (ASI), and spatial specificity of 6609 mall RNAs that are observed with microarrays. The degree of spatial specificity of small RNAs is evaluated by ASI (column F). ASI values are calculated based on mean expression values of the terminal timepoint (day 10) of the three lineages that are corresponding to three tissues, namely, immature hepatocyte (IH), metanephric mesenchyme (MM), and neural progenitors (NPC). The mean expression values of IH, MM, and NPC are shown in column C, D, and E, respectively. The spatial specificity of a small RNA is established as the tissue with the highest expression values among IH, MM, and NPC (column G). Small RNAs are sorted with decreasing ASI values from top to bottom.

**Table S4. Spatiotemporal-specific small RNA indicated by ASI and DE.** Listing of 1318 real spatiotemporal-specific small RNAs with both spatial specificity (from ASI analysis) and temporal specificity (absolute fold-change > 1 and post hoc test FDR-value < 0.05 between any of two timepoints). Column A to E show the probe ID, small RNAs name, small RNA type, spatiotemporal specificity, and ASI. Results of one-way ANOVA *P*-value, fold-change value and post hoc test FDR*-*value are generated from the comparison of expression values between any of two timepoints. These results are listed from column F to DN. IH-specific small RNAs (yellow background), MM-specific small RNAs (blue background), and NPC-specific small RNAs (red background) are grouped together according to their spatiotemporal specificity.

**Table S5. Spatiotemporal specificity of miRNA families.** Summary of spatiotemporal distribution of 37 miRNA families that contain at least five family members. Column A, B, and C show the family name (labelled in red), total number and name of family members of each miRNA family. MiRNA families are sorted with a decreasing number of family members from top to bottom. All family members are grouped together according to their families. IH-specific miRNAs and their ASI are listed in column D and E. The number and mean ASI of IH-specific miRNA are calculated based on column D and E and listed in column F and G. Similarly, the tissue distribution of MM-specific small RNAs inside each family are listed from column I to L. The tissue distribution of NPC-specific small RNAs inside each family are listed from column N to Q. Results associated with IH, MM, and NPC are labelled in the yellow, blue, and red background, respectively.

**Table S6. Spatiotemporal-specific miRNAs indicated by ASI and their downstream targets.** Listing of 618 real spatiotemporal-specific miRNAs with spatial specificity (from ASI analysis) and temporal specificity (absolute fold-change > 1 and post hoc test FDR*-*value < 0.05 between day 0 and day 10). The spatiotemporal-specific miRNAs are classified into 6 groups, corresponding to miRNAs being upregulated at HD 10, KD 10, and ND 10 (fold change > 1 when compared day 10 to day 0) or miRNAs being downregulated at HD 10, KD 10, and ND 10 (fold change < −1 when compared day 10 to day 0). Column A, C, E, G, I, and K show six groups of spatiotemporal-specific miRNAs. Correspondingly, column B, D, F, H, J, and L show downstream target genes of each group of miRNAs that are identified *in silico* using miRTarBase.

**Table S7. GO enrichment analysis.** Listing of all significant biological processes associated with six groups of downstream targets of spatiotemporal-specific miRNAs. Column A show gene ontology (GO) terms of biological processes associated with each group. Column B, C, and D show the number of genes in the reference list, the actual number of genes in each group (uploading list), and the expected number of genes in each group (uploading list) related to each GO term, respectively. Column E to H show the over or under status (compared to 1) according to fold enrichment, fold enrichment, raw *P*-value, and FDR, respectively. The GO terms are grouped together according to their associated gene lists. Within each group, GO terms are sorted with decreasing fold enrichment from top to bottom.

